# Neurometabolite Mapping Highlights Elevated Myo-inositol Profiles within the Developing Brain in Down Syndrome

**DOI:** 10.1101/2020.07.20.211805

**Authors:** Prachi A. Patkee, Ana A. Baburamani, Katherine R. Long, Ralica Dimitrova, Judit Ciarrusta, Joanna Allsop, Emer Hughes, Johanna Kangas, Grainne McAlonan, Mary A. Rutherford

## Abstract

The neurodevelopmental phenotype in Down Syndrome (DS), or Trisomy 21, is variable including a wide spectrum of cognitive impairment and a high risk of early-onset Alzheimer’s disease (AD). A key metabolite of interest within the brain in DS is Myo-inositol (Myo-ins). The NA+/Myo-ins co-transporter, is located on human chromosome 21 and is overexpressed in DS. In adults with DS, elevated brain Myo-ins has previously been associated with cognitive impairment and proposed as a risk marker for progression to AD. However, it is unknown if brain Myo-ins is increased earlier in development.

The aim of this study was to assess Myo-ins and key brain metabolites [N-acetylaspartate (NAA), Choline (Cho) and Creatine(Cr)] in the developing brain in DS and aged-matched controls. To achieve this we used mass spectrometry in early (10-20 weeks post conception) *ex vivo* fetal brain tissue samples from DS (n=14) and control (n=30) cases; and *in vivo* magnetic resonance spectroscopy (MRS) in neonates with DS (n=18) and aged matched controls (n= 25) scanned just after birth (36-45 weeks postmenstrual age.

We observed elevated Myo-ins in the *ex vivo* fetal cortical brain tissue in DS compared with controls. Relative to reference metabolites Cho and Cr, we also detected elevated ratios of Myo-ins and NAA *in vivo* in the basal ganglia and thalami, in neonates with DS, when compared to age-matched typically developing controls. Thus, a higher level of brain Myo-ins was evident as early as 10 weeks post conception and was measurable *in vivo* from 36 weeks post-menstrual age. Future work will determine if this early difference in metabolites is linked to cognitive outcomes in childhood or has utility as a potential treatment biomarker for early intervention.

## Introduction

Down Syndrome (DS) is a complex genetic condition, resulting from the triplication of human chromosome 21 (HSA21; (Trisomy 21), and is associated with a wide spectrum of neurodevelopmental outcomes (Karmiloff-Smith *et al.*, 2016; Baburamani *et al.*, 2019). Individuals typically present with cognitive deficits, and variable impairments in speech, language and motor functions, coupled with a high risk of early-onset dementia and Alzheimer’s disease (AD) (Rachidi and Lopes, 2007; Wiseman *et al.*, 2015; Karmiloff-Smith *et al.*, 2016). There is limited understanding of brain development in DS, but early alterations may contribute to the variations in lifelong neurodevelopmental outcomes and ultimately to the risk for early-onset AD. Recent *in vivo* studies have demonstrated altered structural brain development in DS from as early as 20 weeks gestation, but it is not known whether these are associated with abnormal metabolic profiles (Patkee *et al.*, 2020). Metabolic derangements, specifically with an elevation of myo-inositol (myo-ins), have been described across development in mouse models of DS, as well as in the child and adult human brain with DS, but there have been no studies assessing brain metabolism at much younger ages.

Myo-ins is of particular interest as the NA+/Myo-inositol co-transporter (SLC5A3) is located on HSA21 and is overexpressed in DS (Berry, Wang and Dreha, 1999). Magnetic resonance spectroscopy (MRS) studies in the adult DS population have reported significantly elevated ratios of myo-ins in DS brains, with and without the neuropathological hallmarks of AD, as compared with healthy adult controls (Lin *et al.*, 2016). Higher levels of Myo-ins have been found to correlate with poor cognitive performance, suggesting that this metabolite might have utility for monitoring response to treatment and/or constitute a treatment target (Lin *et al.*, 2016). Consistent with the latter, studies in mouse models of DS have shown that higher levels of Myo-ins could be reduced by treatment with lithium, which also rescued both synaptic plasticity and memory deficits (Huang *et al.*, 1999; Contestabile *et al.*, 2013). Whilst such studies have made valuable first steps to revealing a link between the metabolite profile of DS and functional outcomes, it is unknown when this alteration in Myo-ins (and/or other metabolites supporting brain function) emerges.

Current therapeutic approaches to improve cognition in DS, target children and adults; and these have had limited success. For example, a clinical trial of lithium in adults with DS did not report efficacy (Murphy, 2011). Arguably however, intervention may be needed at an earlier timepoint. Identification of neural differences much earlier in the life course in DS, could potentially offer opportunities for more effective intervention. We and others have recently demonstrated that fetuses and neonates with DS have reduced whole brain, cortical and cerebellar volumes from as early as 20 weeks gestational age (GA), using *in vivo* volumetric MRI (Tarui *et al.*, 2019; Patkee *et al.*, 2020). Whether such anatomical differences are accompanied by early differences in the metabolite levels, that reflect neuronal and glial function, and alter neurodevelopmental outcome, remains unknown.

The aim of this study was to determine whether the brain metabolite profile and Myo-ins, specifically, is already altered in DS in during early development *in utero*. We used *in vivo* MRS to assess the metabolic profile of brain tissue (basal ganglia and thalami) in neonates with DS compared to typically developing controls. To complement this, we used data from a second study and analysed the metabolite profile in post-mortem fetal cortical brain tissue using Mass Spectrometry. In addition to Myo-ins, other metabolites of interest we measured were N-acetylaspartate (NAA), Choline (Cho) and Creatine (Cr). We hypothesised that there would be significant alterations in metabolic ratios between the DS and control groups, with increases in Myo-ins levels specifically, in the brain of neonates with DS and secondly, increases in absolute Myo-ins at earlier gestations in brain tissue from foetuses with DS compared with controls

## Methods

### Magnetic Resonance Spectroscopy Study

Ethical approval for this study was obtained from the West London and GTAC Research Ethics Committee (REC) for DS participants (07/H0707/105); and from the Dulwich NREC for the control population (12/LO/2017). Informed consent was obtained from the legal guardians of all infants, prior to imaging at St. Thomas’ Hospital, London, UK. All methods were carried out in accordance with the relevant guidelines and regulations.

### Participants

Participants who had previously had a fetal scan (Patkee *et al.*, 2020) and consented during their initial scan to be contacted post-delivery, were invited for a neonatal scan up to 46 weeks post-menstrual age (PMA). Neonates with confirmed DS were also recruited from the neonatal unit or postnatal wards at St Thomas’ Hospital. Participants with DS, some of whom had other non-brain congenital abnormalities such as cardiac defects or gastrointestinal malformations, were included in this study. Clinical details for the neonates with DS can be found in Table 1. Control group neonates were recruited from South London and South East of England antenatal centres as part of the Brain Imaging in Babies Study (BIBS) and scanned at the Centre for the Developing Brain (St. Thomas’ Hospital, London, UK). All infants were scanned on the same scanner with identical imaging protocols. Neonates were included as controls if they had a normal brain appearance on the neonatal MRI, with no other congenital or chromosomal abnormalities, and an uneventful delivery, as acute birth-related events could potentially change metabolite composition even in the presence of normal MR imaging. Additionally, all control neonates had no known immediate family members with any neurodevelopmental or behavioural conditions (e.g. autism and ADHD). Previous MRS studies in the paediatric population found statistically significant differences between DS and control groups with 20 participants in each cohort (Śmigielska-Kuzia *et al.*, 2010).

**Table 1:** Age at scan and clinical characteristics of neonates with DS. Gestational age (GA), post-menstrual age (PMA), atrioventricular septal defect (AVSD), atrial septal defect (ASD), ventricular septal defect (VSD) and patent ductus arteriosus (PDA)

| Clinical details of the DS cohort |  |  |  |  |  |
| --- | --- | --- | --- | --- | --- |
| DS Subject | Age at birth (GA weeks) | Neonatal scan (PMA weeks) | Sex (Male/Female) | Cardiac abnormality | Other medical conditions |
| 1 | 38.43 | 41.29 | F | VSD | x |
| 2 | 36.43 | 43.00 | F | AVSD | x |
| 3 | 36.57 | 39.00 | M | x | Duodenal atresia |
| 4 | 37.57 | 40.57 | F | x | Duodenal atresia |
| 5 | 32.29 | 36.14 | F | VSD | Duodenal atresia |
| 6 | 38.14 | 41.43 | F | AVSD and PDA | Duodenal atresia |
| 7 | 41.71 | 43.00 | M | ASD | Hirschsprung's disease |
| 8 | 38.86 | 45.57 | F | Tetralogy of Fallot | Hirschsprung's disease |
| 9 | 36.43 | 38.43 | F | Tetralogy of Fallot, PDA | x |
| 10 | 37.71 | 43.57 | F | AVSD, small left ventricle, hypoplastic arch | x |
| 11 | 38.43 | 39.71 | M | AVSD and hypoplastic aortic arch | x |
| 12 | 36.71 | 44.57 | M | x | x |
| 13 | 38.71 | 44.71 | F | VSD and PDA | Congenital hypothyroidism |
| 14 | 37.00 | 37.57 | M | AVSD and coarctation of the aorta | x |
| 15 | 35.29 | 38.86 | M | ASD | Hirschsprung's disease |
| 16 | 37.00 | 37.86 | F | AVSD | x |
| 17 | 37.14 | 38.00 | M | x | x |
| 18 | 39.43 | 39.43 | M | AVSD | x |

### Neonatal Image Acquisition

MR scanning of the DS and control neonates was performed on a Philips Achieva 3-Tesla system (Best, The Netherlands) at the Centre for the Developing Brain (St. Thomas’ Hospital, London, UK) using a dedicated 32 channel neonatal head coil (Hughes *et al.*, 2017). Infants were placed in supine and secured within the scanning shell and a neonatal positioning device was used to stabilise the head to reduce movement (Hughes *et al.*, 2017). Auditory protection was comprised of earplugs moulded from silicone-based putty placed in the outer ear (President Putty, Coltene/Whaledent Inc., OH, USA), neonatal earmuffs over the ear (MiniMuffs, Natus Medical Inc., CA, USA) and an acoustic hood placed over the scanning shell. Sedation was not administered, and all babies were scanned during natural sleep, following a feed. An experienced neonatologist or neonatal nurse was present during the examination and pulse oximetry, heart rate, and temperature was monitored throughout the scan.

T2-weighted images were acquired in the coronal, sagittal and transverse planes using a multi-slice turbo spin echo sequence. Two stacks of 2D slices were acquired using the scanning parameters: TR = 12 s; TE = 156 ms; slice thickness = 1.6 mm with a slice overlap = 0.8 mm; flip angle = 90 degrees; matrix /field of view= 320 x 320 x 125; and an in-plane resolution: 0.8×0.8 mm. All T2-weighted images were reviewed to assess image quality and those with excessive motion and uninterpretable spectra were excluded. Visual analysis of all neonatal MR images was performed by a specialist perinatal radiologist to exclude additional anomalies and confirm appropriate appearance for gestation. Image datasets showing an overt additional structural malformation or brain injury not considered to be part of the expected DS phenotype were excluded. The data regarding T2-weighted volumetric findings are detailed in Patkee et. al., (2020).

### Spectral Acquisition

A PRESS sequence was used to obtain the MR spectra in the neonatal brains. A region of interest 20 x 20 x 20 mm^3^ was placed centrally over the left basal ganglia and thalami to acquire spectroscopy (Fig 1). Previous studies on early brain development have observed higher individual metabolite levels in the deep grey matter, compared to other regions such as frontal lobe white matter (Kim, Barkovich and Vigneron, 2006; Tomiyasu *et al.*, 2013; Lally *et al.*, 2019). This is additionally beneficial due to the reduced likelihood of contamination by scalp fat. MRS uses a double spin echo sequence (90^°^ – 180^°^ – 180^°^) to improve signal quantity.

**Figure 1:**
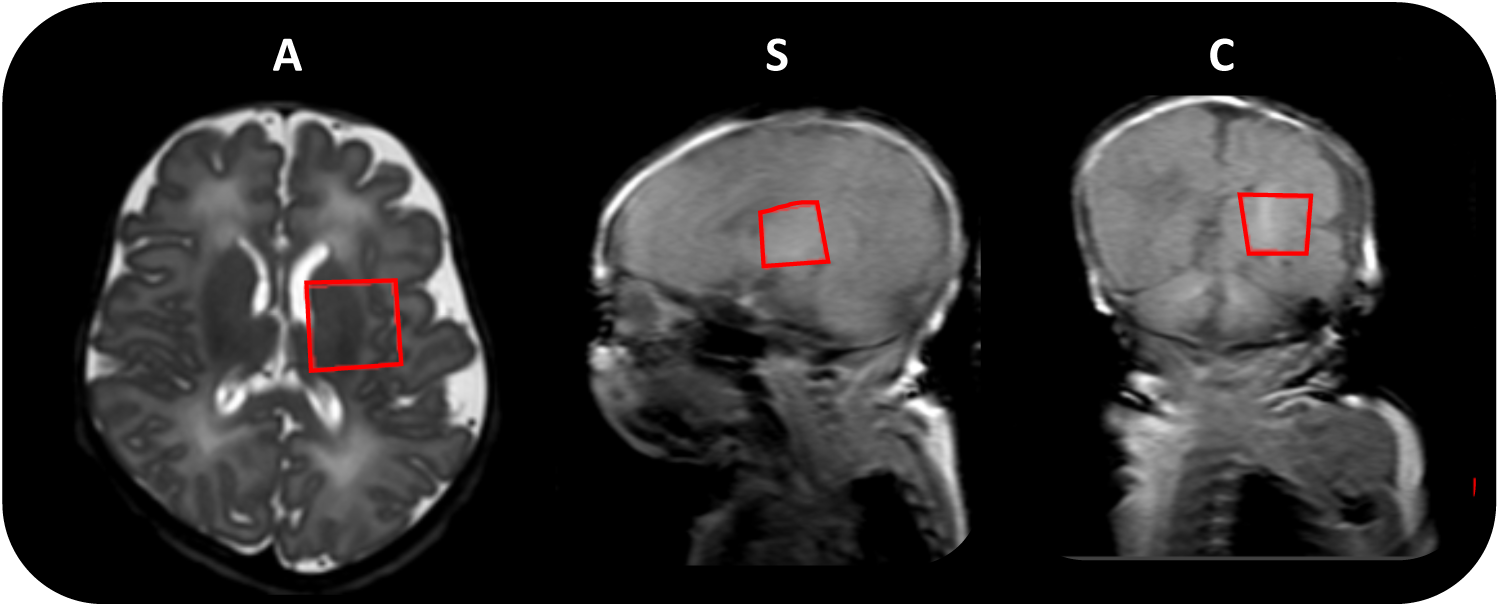
Neonatal MRS acquisition. Region of interest size 20 x 20 x 20 mm^3^ (left basal ganglia and thalami) indicated by the red box, in the axial (A), sagittal (S) and coronal (C) planes respectively.

Each spectrum took approximately 3 minutes to obtain and was acquired in blocks of four averages (producing 32 individual spectra) and saved independently, allowing for the elimination of individual corrupted spectra as a result of excessive motion or contamination of peaks, prior to post-processing the data. Water suppression was performed by selective excitation. This is necessary to avoid metabolite signals being overwhelmed by the large water peak and to ensure that the baseline of the spectrum is not distorted, which can make it difficult to quantify individual smaller peaks. In this study, metabolites of interest were measured at two echo times, TE 55 and 144 ms.

### Processing and Analysis of Spectral Data

The software installed on the Philips MRI scanner performs some basic rapid automatic processing for immediate quality assessment of the acquired spectrum. For a complete quantitative analysis of metabolite ratios, spectra were post-processed offline using the jMRUI software package (http://www.jmrui.eu/, version 3.0. The spectral analysis aims to estimate signal intensities or areas under the peaks corresponding to the metabolites. Individual spectra were quality assessed for high SNR by the height of metabolite peaks as compared to background noise. Data with significant motion or those with aberrant water peaks were discarded. The remaining spectra with a residual water peak at the accepted threshold were summed together to produce a final spectrum, after which the water peak at 4.7ppm was removed (Fig 2). In this study we focused on the following metabolites of interest (resonant frequency); N-acetylaspartate (2.0 ppm), Creatine (3.0 ppm), Choline (3.2 ppm) and Myo-inositol (3.5 ppm).

**Figure 2:**
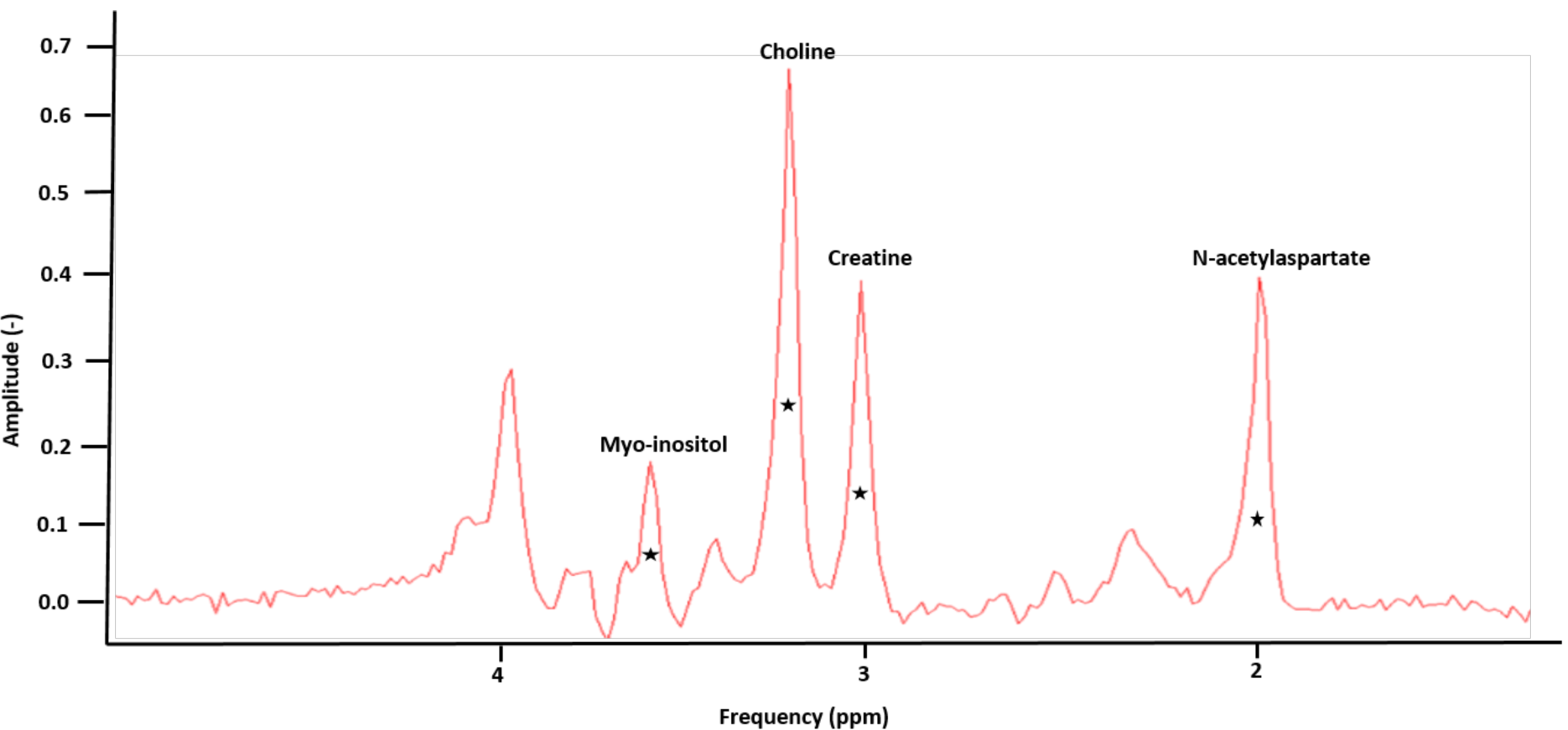
MR spectrum from a neonate with DS scanned at 38 weeks PMA, using a PRESS sequence at an echo time of 144ms. Peaks for Myo-inositol (3.5 ppm), Choline (3.2 ppm), Creatine (3.0 ppm) and N-acetylaspartate (2.0 ppm), are labelled, following the removal of the H_2_O peak.

### Quantification of Metabolites

The jMRUI software allows for the time-domain analysis of MR spectra using the AMARES programme. Expected peaks for each of the metabolites were identified on the spectrum based on their frequency (ppm). The top and midpoint of each peak were identified and based on this prior knowledge a curve was fit by the software for each metabolite. Once the curves had been fitted, metabolites ratios were calculated for the two echo times as follows: TE 55 ms; Myo:Cho, Myo:Cr, Cho:Cr and TE 144 ms; NAA:Cho, NAA:Cr, Cho:Cr.

The different peaks in the spectrum represent individual metabolites and are identified by the frequency at which they resonate, represented in parts per million (ppm). In MRS, the total area under a metabolite resonance in an MR spectrum is directly proportional to the concentration of the metabolite. As the MR signal is affected by factors such as relaxation time and the acquisition method, amongst others, it is not possible to establish an absolute concentration without accounting for these confounding elements. At present, metabolites are usually calculated as a relative quantification and therefore are expressed as ratios rather than as absolute concentrations. This is particularly useful when detecting subtle or more sensitive changes and can be more accurate than an absolute quantification; this is true in cases with pathology where one metabolite concentration rises whilst the other declines (Jansen *et al.*, 2006). All neonatal spectra were processed by the same operator.

## Mass Spectrometry Study

### Fetal Brain Tissue Samples

Human fetal cortex material was provided by the Joint MRC/Wellcome Trust (MR/R006237/1) Human Developmental Biology Resource (HDBR) (www.hdbr.org). HDBR provided fresh fetal cortex tissue that was snap-frozen for DS (n=14) and age-matched controls (n=30) in the fetal age range 10 to 20 post-conception weeks (PCW). Genetic diagnosis confirmed presence of Trisomy 21 (DS). There were no other known abnormalities in either group. HDBR specific ethics approval (18/LO/0822, 18/NE/0290) was sought and tissue was collected following termination of pregnancy, following Human Tissue Act (HTA) regulations. Cortical tissue was then selected, snap frozen and stored at -80 ^°^C. Cases that were used are detailed in Table 2.

**Table 2:** Case details of fetal brain samples with DS and aged-matched controls. Age at birth, in post-conceptional weeks (PCW), sex and other genetic/co-morbidities are listed.

| Subject ID | Age (pcw) | Sex (Male/ Female) |
| --- | --- | --- |
| DS 1 | 10 | F |
| DS 2 | 11 | Unknown |
| DS 2 | 12 | M |
| DS 4 | 12 | F |
| DS 5 | 12 | M |
| DS 6 | 12 | M |
| DS 8 | 13 | F |
| DS 9 | 13 | M |
| DS 10 | 14 | M |
| DS 11 | 14 | M |
| DS 12 | 14 | M |
| DS 13 | 15 | F |
| DS 14 | 17 | M |
| DS 15 | 19 | M |
| Control 1 | 10 | F |
| Control 2 | 10 | F |
| Control 3 | 11 | M |
| Control 4 | 11 | M |
| Control 5 | 11 | M |
| Control 6 | 12 | F |
| Control 7 | 12 | M |
| Control 8 | 12 | M |
| Control 9 | 13 | M |
| Control 10 | 13 | M |
| Control 11 | 13 | M |
| Control 12 | 14 | F |
| Control 13 | 14 | F |
| Control 14 | 14 | M |
| Control 15 | 15 | M |
| Control 16 | 15 | F |
| Control 17 | 16 | F |
| Control 18 | 16 | M |
| Control 19 | 16 | M |
| Control 20 | 17 | F |
| Control 21 | 17 | F |
| Control 22 | 17 | M |
| Control 23 | 18 | M |
| Control 24 | 18 | F |
| Control 25 | 19 | F |
| Control 26 | 19 | M |
| Control 27 | 19 | F |
| Control 28 | 20 | M |
| Control 29 | 20 | F |
| Control 30 | 20 | F |

### Sample preparation

Human fetal cortex tissue samples were stored at –80°C. The cortical tissue used included a variety of regions, as due to the nature of these fetal samples it is often difficult to identify the exact cortical region tissue originates from across the gestational ages. To ensure this did not affect the results, cortical tissue was randomly assigned for the mass spectrometry analysis in both DS and control groups. Approximately 0.1g of brain tissue from the cortex was collected on dry ice and weighed. 1 ml of HPLC grade Methanol (Sigma, UK) was added and samples were then homogenised for 20 minutes at 30 Hz using TissueLyser II device and centrifuged at 14,000 rpm for 0.5 minute (longer if required). Clear supernatants were then transferred to a new tube on dry ice, samples were then stored at -80 °C until analysis. Analysis of Myo-Inositol and NAA was performed with Gas Chromatography Mass Spectrometry (GCMS) and for Creatine and Choline using Liquid Chromatography Mass Spectrometry (LCMS). All metabolite values for an individual fetal sample were measured from a single tissue sample to eliminate inter-assay variability. All analysis was run by the Mass Spectrometry Core Facility, Kings College London (KCL, UK).

### Gas Chromatography Mass Spectrometry (GCMS) for Myo-Inositol and NAA

Samples (10ul; Blank, Calibration standard or test samples) were added to tubes containing internal standard-200 (20 µl) + solution (50µl) (mixture of internal standards (200 ug/ml Myo-inositol internal standard, 50 ug/ml NAA internal standard solution). These were evaporated to nearly dry under N2 at 40±5°C. Pyridine (10 µl) and BSTFA (40 µl) were then added and the samples were vortexed, heated at 80±5°C for 30 minutes, cooled down to room temperature and transferred to injection vials. Calibration ranges were 50-2000 µg/ml for Myo-Inositol (Cat# M01914, Fluorochem Limited, UK; Internal Standard Myo-inositol-D6 Cat# 1665997, Toronto Research Chemicals, Canada) and 1-100 µg/ml for NAA (Cat# 00920, SLS, UK). Samples were analysed on an Agilent GCMSD system. GC-MS parameters are detailed in Table 3.

**Table 3:**
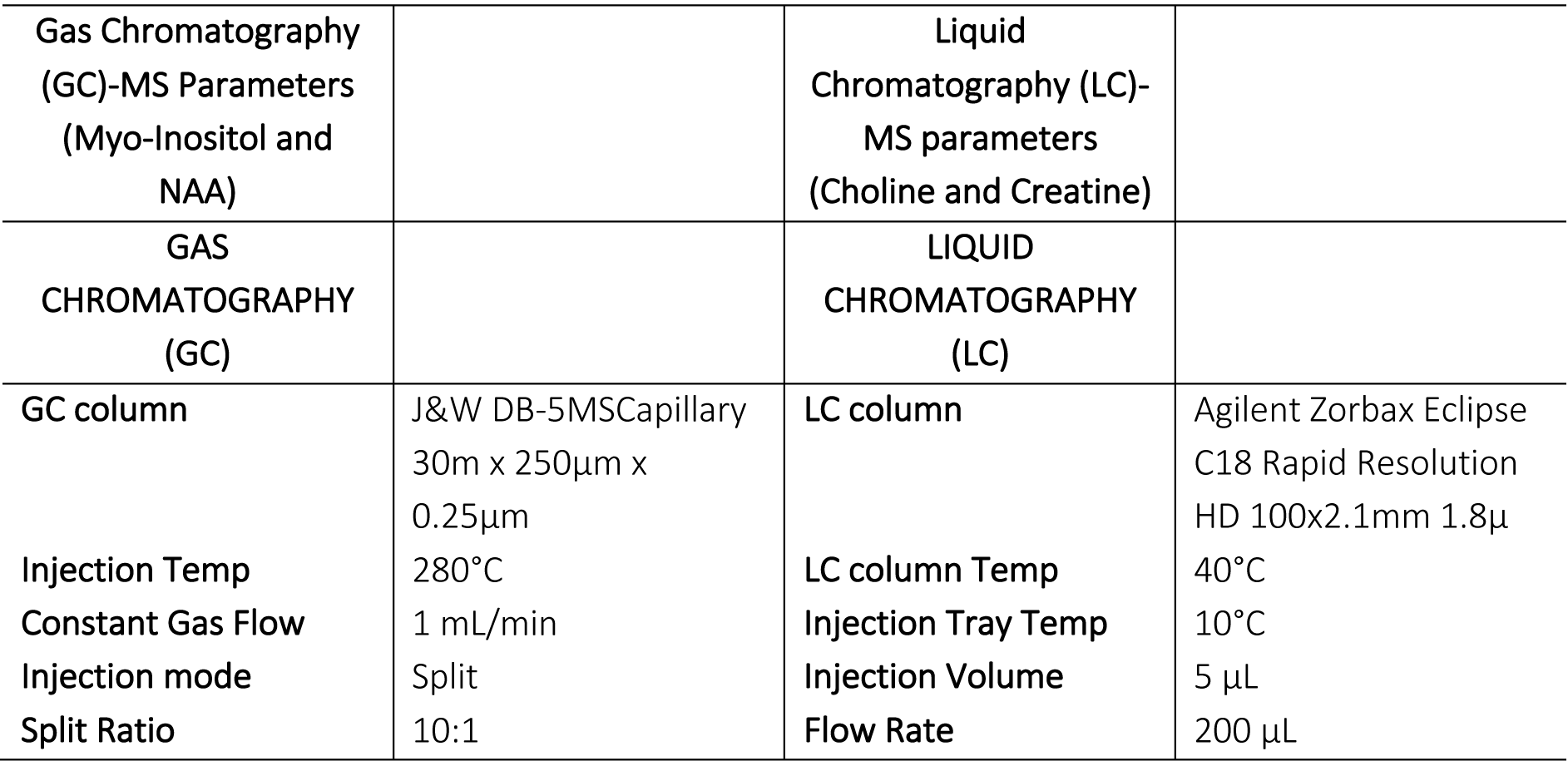
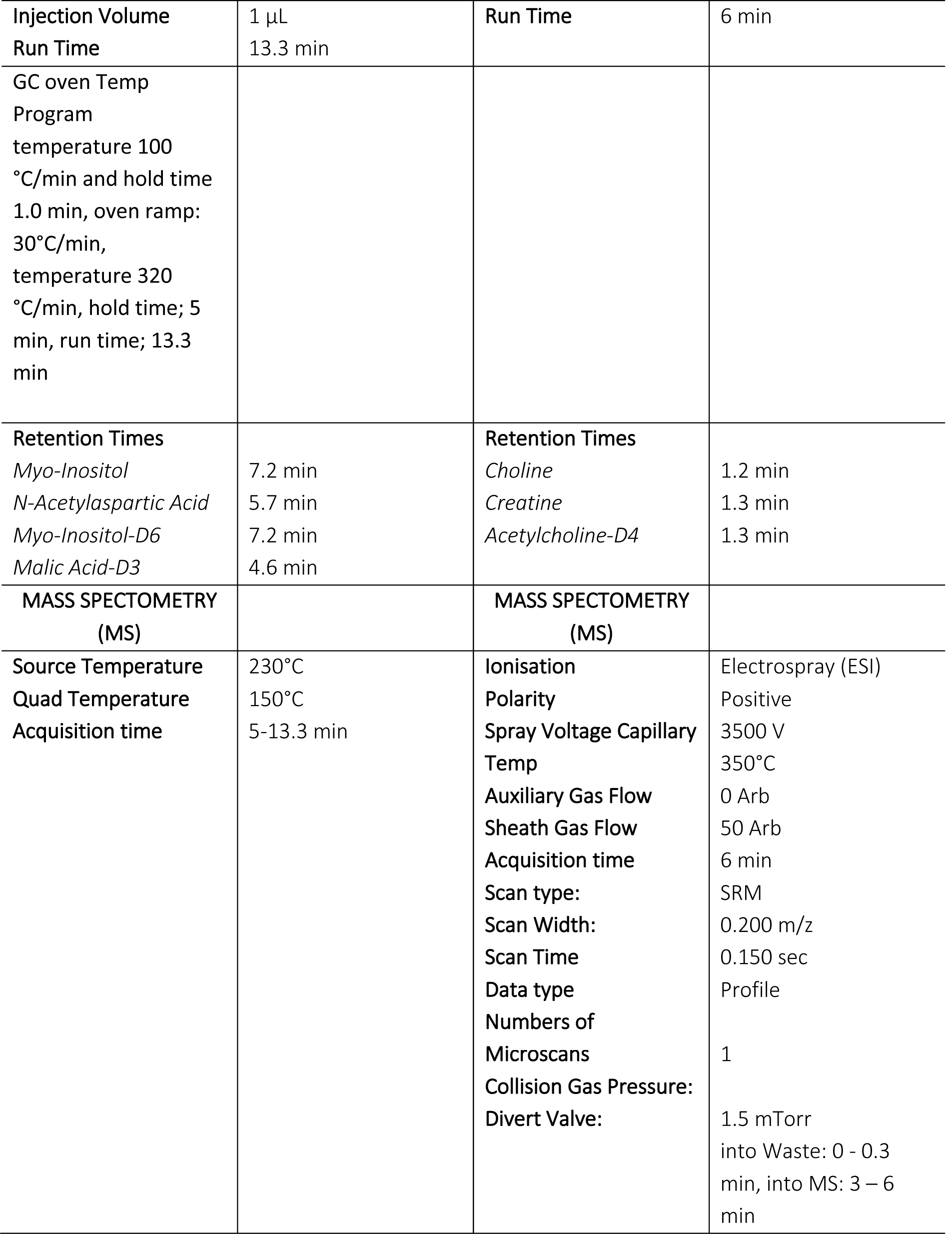
GCMS and LCMS parameters.

| Gas Chromatography (GC)-MS Parameters (Myo-Inositol and NAA) |  | Liquid Chromatography (LC)-MS parameters (Choline and Creatine) |  |
| --- | --- | --- | --- |
| GAS CHROMATOGRAPHY (GC) |  | LIQUID CHROMATOGRAPHY (LC) |  |
| GC column | J&W DB-5MSCapillary 30m x 250μm x 0.25μm | LC column | Agilent Zorbax Eclipse C18 Rapid Resolution HD 100x2.1mm 1.8μ |
| Injection Temp | 280°C | LC column Temp | 40°C |
| Constant Gas Flow | 1 mL/min | Injection Tray Temp | 10°C |
| Injection mode | Split | Injection Volume | 5 μL |
| Split Ratio | 10:1 | Flow Rate | 200 μL |
| Injection Volume | 1 µL | Run Time | 6 min |
| Run Time | 13.3 min |  |  |
| GC oven Temp<br>Program<br>temperature 100<br>°C/min and hold time<br>1.0 min, oven ramp:<br>30°C/min,<br>temperature 320<br>°C/min, hold time; 5<br>min, run time; 13.3<br>min |  |  |  |
| Retention Times<br>*Myo-Inositol*<br>*N-Acetylaspartic Acid*<br>*Myo-Inositol-D6*<br>*Malic Acid-D3* | 7.2 min<br>5.7 min<br>7.2 min<br>4.6 min | Retention Times<br>*Choline*<br>*Creatine*<br>*Acetylcholine-D4* | 1.2 min<br>1.3 min<br>1.3 min |
| MASS SPECTROMETRY<br>(MS) |  | MASS SPECTROMETRY<br>(MS) |  |
| Source Temperature<br>Quad Temperature<br>Acquisition time | 230°C<br>150°C<br>5-13.3 min | Ionisation<br>Polarity<br>Spray Voltage Capillary<br>Temp<br>Auxiliary Gas Flow<br>Sheath Gas Flow<br>Acquisition time<br>Scan type:<br>Scan Width:<br>Scan Time<br>Data type<br>Numbers of<br>Microscans<br>Collision Gas Pressure:<br>Divert Valve: | Electrospray (ESI)<br>Positive<br>3500 V<br>350°C<br>0 Arb<br>50 Arb<br>6 min<br>SRM<br>0.200 m/z<br>0.150 sec<br>Profile<br>1<br>1.5 mTorr<br>into Waste: 0 - 0.3<br>min, into MS: 3 – 6<br>min |

### Liquid Chromatography Mass Spectrometry (LCMS) for Creatine and Choline

Samples (100 µl; Blank, Calibration or test samples) were added to individual injection vials containing Internal Standard-2u (50 µl; 2µg/mL Acetylcholine-D4, Cat#D-1555, CDN Isotopes, Canada) and DS1 (900 µl, contains 0.1% Formic Acid in 50 % HPLC grade Acetonitrile), vortexed, and injected into the LCMS system. DS1 was run between high and low concentration samples and between sets of the test sample. Calibration ranges were 10 – 200 µg/ml for Creatine (Cat# BS-9561E, BioServ UK Limited) and 0.10-35.0 µg/ml for Choline (Cat# 67-48-1, Sigma Aldrich, UK). Samples were analysed on a Thermo Accela Pump and CTC Auto sampler coupled to a Thermo TSQ Quantum Access. LC Solvent A (0.1% Formic Acid in water); LC Solvent B (0.1% Formic Acid in Acetonitrile). LCMS parameters detailed in Table 3.

### Statistical Analysis

Statistical analysis was performed using MATLAB (Release: 2017a, The MathWorks, Inc., Natick, MA, USA), and visualised in python 3.7 (www.python.org). Linear mixed-effects models (LME) were used to compare results between the two groups (DS and controls) in both the mass spectrometry and MRS studies, controlling for the covariate PMA at scan as a fixed effect. Cohen’s *d* values were calculated to quantify effect sizes and the magnitude of interactions (Cohen, 1988; Rosenthal and Rosnow, 2008). Effect sizes were interpreted as small (Cohen’s *d* value ≤ 0.4), medium (0.5-0.7) and large (≥ 0.80) (Cohen, 1988). A post-hoc False Discovery Rate (FDR) test was performed for multiple comparisons, using the Benjamini-Hochberg (1995) procedure, controlling the alpha error to 5%.

## Results

### Neonatal Magnetic Resonance Spectroscopy

A total of 21 neonates with DS were scanned (Patkee *et al.*, 2020), 18 had usable and interpretable spectral acquisitions. For comparison, 25 control neonates were included as part of this study. Of these, 21 neonates had interpretable spectral acquisitions (Table 4).

**Table 4:** Summary of the DS and Control neonates with interpretable spectra. Gestational age (GA) and post-menstrual age (PMA).

|  | Down Syndrome neonates | Control neonates |
| --- | --- | --- |
| Total number (n) | 18 | 25 |
| GA at birth, median (range) weeks | 37.00, (32.29-41.71) | 40.00, (36.57-41.86) |
| PMA at scan, median (range) weeks | 40.14, (36.14-45.57) | 42.86, (39.29-44.29) |
| Males/Females (n) | 8/10 | 16/9 |
| MRS acquired (TE 55ms) (n) | 10 | 21 |
| MRS acquired (TE 144ms) (n) | 16 | 23 |

The Myo:Cho and Myo:Cr ratios in the control cohort appeared to decrease with advancing PMA at scan. There was marked variability in Myo-ins ratios across PMA in the DS population, with no obvious decrease with PMA at scan. Myo:Cho and Myo:Cr ratios were elevated in DS neonates compared to controls, corrected for PMA at scan, with group differences of large effect sizes (*p* < 0.05, Table 5, Fig 3A, 3B). NAA:Cho and NAA:Cr ratios were found to increase with advancing PMA in both the DS and control groups. There were increased ratios of NAA:Cho and NAA:Cr in the DS cohort compared with controls (*p* < 0.05, Table 5, Fig 3C, 3D). Cho:Cr ratios decreased with increasing PMA in both cohorts. The significance of Cho:Cr ratios in the DS and control neonates varied based on the TE of the acquisition. The Cho:Cr (TE 55 ms) ratio, was not to differ between the two groups (*p* =0.12); however, Cho:Cr (TE 144 ms) ratio was lowered in DS neonates (*p* < 0.05).

**Table 5:** LME results, Cohen’s *d* values and effect sizes for the DS vs control cohort - MRS data.

| **Metabolite Ratios** | **Estimate** | **Standard Error** | **df** | **t-value** | **p-value** | **Cohen's $d$** |
| --- | --- | --- | --- | --- | --- | --- |
| Myo:Cho | -0.31 | 0.05 | 28 | -6.85 | 0.00* | -2.6 |
| Myo:Cr | -0.3 | 0.05 | 28 | -6.72 | 0.00* | -2.5 |
| NAA:Cho | -0.16 | 0.03 | 36 | -5.79 | 0.00* | -1.9 |
| NAA:Cr | -0.1 | 0.05 | 36 | -2.11 | 0.05* | -0.7 |
| Cho:Cr (TE 55ms) | 0.08 | 0.05 | 28 | 1.6 | 0.12 | 0.6 |
| Cho:Cr (TE 144ms) | 0.27 | 0.07 | 36 | 3.91 | 0.00* | 1.3 |

\*= significant result, $p < 0.05$ . Effect sizes were interpreted as small (Cohen's $d$ value $\leq 0.4$ ), medium (0.5–0.7) and large ( $\geq 0.8$ ) (Cohen, 1988). N- acetylaspartate (NAA - 2.0 ppm), Creatine (Cr - 3.0 ppm), Choline (Cho - 3.2 ppm) and Myo-inositol (Myo-ins - 3.5 ppm).

**Figure 3:**
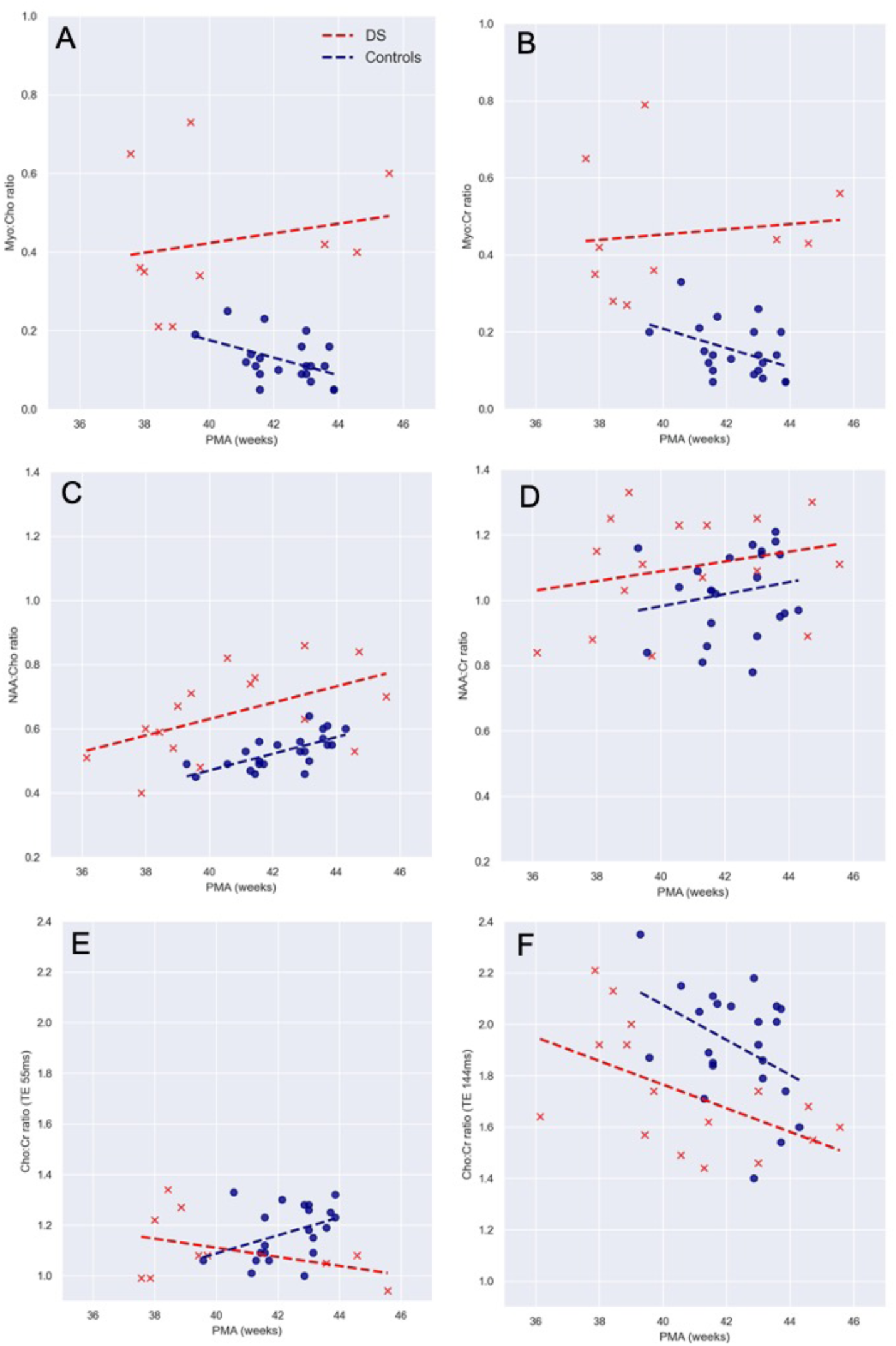
Metabolite ratios in neonates with DS (red circles) and age-matched normal controls (black squares). (A) Myo-inositol:Choline (TE 55ms), (B) Myo-inositol:Creatine (TE 55ms), (C) N-acetylaspartate:Choline (TE 144ms), (D) N-acetylaspartate:Creatine (TE 144ms), (E) Choline:Creatine (TE 55ms), and (F) Choline:Creatine (TE 144ms). Post-menstrual age (PMA).

No significant differences were found in metabolite ratios between male and female neonates in either the control or DS groups. We could not analyse the differences in metabolite ratios in DS participants with (n=14) and without (n=4) a congenital heart defect (CHD), due to insufficient numbers for comparison. Neither, DS neonates without a cardiac defect (n=4), nor those born prematurely (prior to 37 weeks, n=6) were outliers.

### Fetal Mass spectrometry study

We measured the total amount of the metabolites Creatine, Choline, Myo-inositol and NAA (Figure 4; Table 6) in cortical tissue from fetal brain. 14 fetuses with DS (median (range) - 13.50 (10-19) PCW and 30 control fetuses (15.00 PCW, (10-20) PCW) were used in this study. We found a statistically significant elevation in total Myo-inositol in DS fetal brains compared to controls (p < 0.05, d = -0.7), when controlled for age (Figure 4C). This difference appears to be greater in the 10-12 pcw samples analysed. No group differences were observed in Creatine, Choline and NAA. However, the trajectories over gestation appear to be lower in DS as compared to controls. Specifically, Myo-inositol and Creatine appears to be on downward trajectories in DS, as compared to the control population. For comparison with the MRS study, ratios were calculated for the mass spectrometry cohort, from the absolute values (Supp. table 1). The ratios for Myo:Cho and Myo:Cr were found to be significantly different between the two groups (p<0.05), supporting our MRS findings. However, a statistically significant difference was not detected for the other ratios.

**Table 6:** LME results, Cohen’s *d* values and effect sizes for the DS vs control cohort – Mass spec data

| Metabolites (ug/g) | Estimate | Standard Error | df | t value | p value | Cohen's d |
| --- | --- | --- | --- | --- | --- | --- |
| Myo-ins | -370.8 | 170.37 | 41 | -2.18 | 0.04* | -0.7 |
| NAA | 8.15 | 7.03 | 41 | 1.16 | 0.25 | 0.4 |
| Choline | 7.81 | 5.31 | 41 | 1.47 | 0.15 | 0.5 |
| Creatine | 106.07 | 64.83 | 41 | 1.64 | 0.11 | 0.5 |

\*= significant result, $p < 0.05$ . Effect sizes were interpreted as small (Cohen's *d* value $\leq 0.4$ ), medium (0.5–0.7) and large ( $\geq 0.8$ ) (Cohen, 1988). N- acetylaspartate (NAA), Creatine, Choline and Myo-inositol.

**Figure 4:**
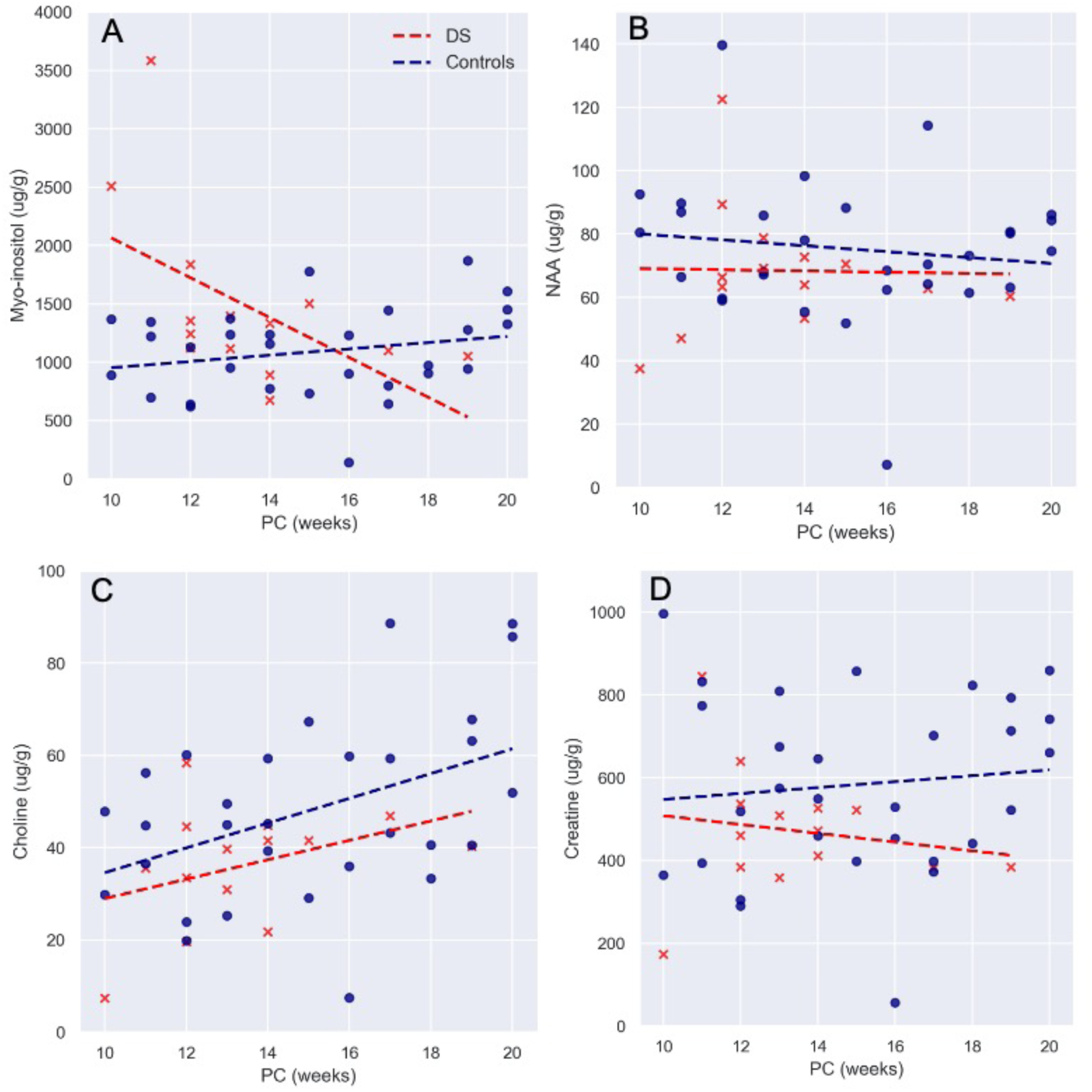
Absolute values for Creatine, Choline, Myo-inositol and NAA in fetal brain tissue for DS (red circles) and age-matched control (black circles) across gestation. Post-conception (PC). Lines indicate mean.

## Discussion

To the best of our knowledge, this is the first study to report elevations in both, absolute total myo-inositol values *ex vivo* in the human fetal brain with DS, and *in vivo* MRS brain data in neonates with DS. In this study, we investigated metabolic profiles in the developing brain with DS, with focus on Myo-ins. We found significant alterations in the metabolite ratios of Myo-ins and NAA in the brain of neonates with DS compared to typically developing neonates. We also found that absolute Myo-ins was elevated in fetal brain tissue in DS, prior to 20 weeks when compared to controls, markedly so earlier in gestation at 10 weeks.

Our study found Myo-ins ratios were significantly higher in the DS neonates, compared to the control population in the deep gray matter. In support of an elevation of Myo-ins in DS, our second study assessing fetal brain tissue with mass spectrometry also demonstrated elevated Myo-ins levels in the cortex, when compared to controls prior to 20 weeks gestation, particularly in the earliest ages examined, 10-12 pcw. Thus far, increased absolute Myo-ins levels have not been reported in the human developing fetal DS brain using mass spectrometry. As the early imbalances in Myo-inositol levels, from 10-12 pcw, may account for the overall observed increase in DS cases, a larger sample size would be needed to determine if there is a consistent, comparative elevation in absolute myo-ins concentration in DS fetuses.

Myo-ins is specifically relevant in the pathology of DS, as the NA+/Myo-inositol co-transporter (SLC5A3) is located on HSA21 (Berry, Wang and Dreha, 1999). Inositol transporters such as sodium-Myoinositol cotransporter-1 (SMIT1), which are encoded by the SLC5A3 gene, are responsible for regulating brain Myo-ins levels, by co-transporting two sodium ions along the concentration gradient to aid active transport of Myo-ins across cell membranes (Fenili *et al.*, 2011; Dai *et al.*, 2016). Myo-ins has been shown to be increased in trisomic cortical neurons (40% compared to diploid), generated from a mouse model trisomic for mouse chromosome 16 (which shows large synteny with HSA21) (Acevedo *et al.*, 1997). Elevated levels of Myo-ins have also been found in amniotic fluid of human DS pregnancies (15-18 weeks GA) (Santamaria *et al.*, 2014). SMIT1 is overexpressed in human DS and associated with increased intracellular levels of Myo-ins in adults (Shonk and Ross, 1995; Cárdenas *et al.*, 2017). Given the association of Myo-ins transporters and HSA21, our results, likely reflect an elevation in incorporated intracellular Myo-ins concentration.

Myo-ins is a glial cell marker and increased levels have been associated with neuroinflammation (Chang *et al.*, 2013). It is speculated that abnormal Myo-ins metabolism may be associated with the initial cognitive impairment in DS as well as being a predisposing factor to AD. Increased concentration of Myo-ins in AD and mild cognitive impairment (without AD) have been supported by histopathologic findings of demyelination and increased gliosis, which is thought to be related to a combination of disintegration of white matter fibres, microglial activation or astrogliosis (Siger *et al.*, 2009). There is increasing evidence from histological studies from mouse models of AD, supporting a link between elevated concentrations of Myo-ins and astrocytic activation (evidenced by an increase in glial fibrillary acidic protein (GFAP) staining), in response to inflammation (Harris *et al.*, 2015).

Increased Myo-ins:Cr have been identified using MRS, in both grey and white matter, in DS patients (Shonk and Ross, 1995) and non-DS patients with AD and dementia, and found to precede associated decreases in NAA, atrophy, neuronal loss and cognitive impairment (Öz *et al.*, 2014). Additionally, higher hippocampal concentrations of Myo-ins have been reported in DS adults without dementia, compared to healthy controls (Beacher *et al.*, 2005). Dietary intake has been shown to directly influence plasma levels of Myo-ins, but not those in the CSF, which is thought to be a consequence of global dysregulation in metabolism or of increased dosage of a single gene on HSA21 (Holub, 1986; Shetty *et al.*, 1995). Shetty, et al., (1995) placed DS (age: 22-63 years) and control (age: 23-69 years) participants on a low monoaminergic diet for 72 hours, prior to collecting samples from a lumbar puncture. They reported a 30-50% increase in Myo-ins concentrations in the CSF of DS subjects compared to controls (Shetty *et al.*, 1995; Fisher, Novak and Agranoff, 2002). As highlighted, the majority of Myo-ins studies have investigated levels in the adult brain, as such, our findings of elevated concentrations in the fetal brain are particularly interesting for further research and as a target for potential interventions.

The association of elevated Myo-ins levels with cognitive impairment in the adult brain, led to a recent clinical trial which aimed to reduce elevated brain Myo-ins levels in adults with DS with scyllo-inositol but found no apparent behavioural or cognitive improvements (Rafii *et al.*, 2018). The authors attribute the result to small participant numbers, short trial duration and the potential practice effects on exploratory cognitive testing. Mouse studies of DS using the Ts65Dn model, have shown promising results in successfully reducing levels of Myo-ins with lithium treatment in the adult mouse, rescuing both synaptic plasticity and memory deficits (Huang *et al.*, 1999; Contestabile *et al.*, 2013). Lithium has been presented as potential treatment option in children with mood disorders, as well as those with intellectual disorders (Yuan *et al.*, 2018). Clinical trials of lithium in children with intellectual disabilities reported positive effects on cognition with only mild, reversible side effects (Yuan *et al.*, 2018), highlighting a potential avenue of research and prospective treatment for DS children with elevated Myo-ins.

In this MRS study we also observed elevated NAA:Cho and NAA:Cr ratios in DS compared to controls. However, this was not apparent prior to 20 weeks gestation, where we saw no difference in absolute NAA values in the fetal brain. Our MRS study also found increasing NAA:Cho and NAA:Cr ratios, with advancing PMA in both our DS and control cohorts. Previous studies in a typically developing fetal cohort (n=35; 30-41 weeks GA) have shown a similar increase in NAA:Cho and NAA:Cr ratios, increasing GA, coupled with a decrease in Cho:Cr (Kok *et al.*, 2002). NAA is one of the most concentrated molecules in the brain. Increasing levels of NAA are reflective of brain maturation and attributable to dendritic and synaptic development and oligodendroglial proliferation and differentiation (Kato *et al.*, 1997; Battin and Rutherford, 2002). Previous studies in adults with DS have reported lower NAA ratios (Śmigielska-Kuzia *et al.*, 2010; Lin *et al.*, 2016). This is thought to be an early marker of neurodegenerative changes, preceding structural changes as visualised by MRI. This is the first evidence of elevated absolute NAA in the early developing brain in DS. The significance of this result is not yet clear and may reflect abnormal acceleration of development and premature exhaustion as a consequence of potential mitochondrial dysfunction (Coskun and Busciglio, 2012)

In this study Cho:Cr ratios were found to decrease with increasing PMA at scan in both the control and DS cohorts. Total Cho and Cr in the fetal brain was not found to differ prior to 20 weeks in the DS and control tissue. Cho plays a role in lipid membrane synthesis and degradation, and in myelination, and is found in both glia and neurons (Kok *et al.*, 2001; Girard *et al.*, 2006). Adults MRS studies in a healthy ageing population found that Cr levels elevated over time, were thought to be a marker of decreased brain energy metabolism and associated with ageing-related mild cognitive impairment, or dementia in more extreme cases (Ferguson *et al.*, 2002). We observed a significant decrease in Cho:Cr in our DS cohort compared to controls, at TE 144ms, but not at TE 55ms. A potential reason for this may be the limited number of cases at TE 55ms between 40-44 weeks PMA. This finding is unlikely to be associated with advanced myelination and maturation of the DS brain, as previous studies have observed delayed myelination in post-mortem fetuses and neonates (Wisniewski and Schmidt-Sidor, 1989). Given that the results from this study show a decrease in the Cho:Cr ratio, it is reasonable to propose that a decrease in choline may be responsible, but perhaps less likely, for the unexpected increase in NAA:Cho ratio.

To assess whether alterations in metabolite values exist and could be measured in utero, we performed a preliminary fetal MRS study on a small cohort of fetal DS cases (n=11) (Patkee *et al.*, 2020), compared with age-matched controls (n=36). Fetal DS data was acquired at a combination of 1.5T and 3T field strengths, whilst the control data was exclusively acquired on a 1.5T scanner. Data was analysed using the same methodology as the neonatal cohort. Unfortunately, much of the data was found to be heavily motion artefacted and corrupted, with only a few cases which had interpretable spectra, therefore the results were deemed to be inconclusive. This is a limitation of the wider study, as we are only able to comment on fresh-frozen fetal tissue, as *in vivo* fetal MRS is thus far unreliable. Future work will aim to optimise MRS sequences in the fetus to account for motion and to acquire a greater volume of data, to account for the corrupted spectral acquisitions.

### Limitations

Ideally, we would aim to acquire MR spectra at both echo times and from more than one region of interest. However, due to technical challenges, such as excessive motion or a neonate becoming unsettled during a scan, it was not always possible to acquire adequate quality MRS at both echo times in all cases. Shorter TEs yield a much better SNR for the detection of metabolites, however at these TEs lipid signal contamination can be found. Longer TEs produce spectra with a much clearer baseline and a reduction in lipid signals which allows for easier metabolite quantification. Due to the intrinsic variation in T2 relaxation times across the metabolites between short and long TEs, normal spectra heights may/will vary when measured at different TEs. It is evident that both cohorts have small sample sizes for spectral analysis. However, we detected significant differences in the Myo:Cho, Myo:Cr, NAA:Cho, NAA:Cr and Cho:Cr (TE136 ms) ratios between DS and control neonates. These novel changes have not been previously reported in the developing brain this early in development. There are several limitations linked to the nature of working with human fetal tissue samples, that we have taken steps to address in our experimental design. Firstly, whilst a wide range of gestational ages were sampled, wherever possible three individual samples were used per age for the controls. Secondly, in both DS and control the cortical tissue used for the mass spectrometry analysis was selected randomly, to avoid bias in the cortical areas analysed. Thirdly, due to the ages of the samples collected, the majority were collected from surgical, not medical, termination procedures, reducing the effect on tissue metabolites. Finally, once analysed by mass spectrometry, all readings were corrected for cortical tissue weight and DS samples were compared to the age-matched controls.

### Conclusions

In conclusion, we found significantly elevated Myo-Ins in the DS fetal brain, and elevated ratios of Myo:Cho, Myo:Cr, NAA:Cho and NAA:Cr in the developing neonatal brain with DS, compared to a typically developing population. DS is a multifactorial disorder with a wide spectrum of phenotypes attributable to a significant degree of individual variability in genotypes (Karmiloff-Smith *et al.*, 2016; Baburamani *et al.*, 2019). The differences in metabolite ratios may reflect ongoing metabolic deviations that may underlie some of the altered brain volumes previously reported (Patkee *et al.*, 2020) in DS and may be correlated with subsequent cognitive delay and potentially with the development of dementia in later life. Future studies will explore possible correlations with childhood neurodevelopmental assessments in this cohort to ascertain whether metabolite levels predict the degree of early neurocognitive impairment in DS.

## Acknowledgements

We thank the parents and children who participated in this study. The authors gratefully acknowledge staff from the Centre for the Developing Brain at King’s College London and the Neonatal Intensive Care Unit at St. Thomas’ Hospital; particularly the research radiologists, radiographers, clinicians, neonatal nurses, midwives and the administrative teams. In addition, we wish to thank all our obstetric and fetal medicine colleagues from our patient identification sites who have referred participants to us. We would also like to thank Dianne Gerrelli, Steve Lisgo, Berta Crespo and their teams at the HDBR for their invaluable support, and Anna Caldwell, John Halket and their team at the Mass Spectrometry facility at King’s College London for their expertise and support.

## Supplementary materials

**Supplementary table 1:**
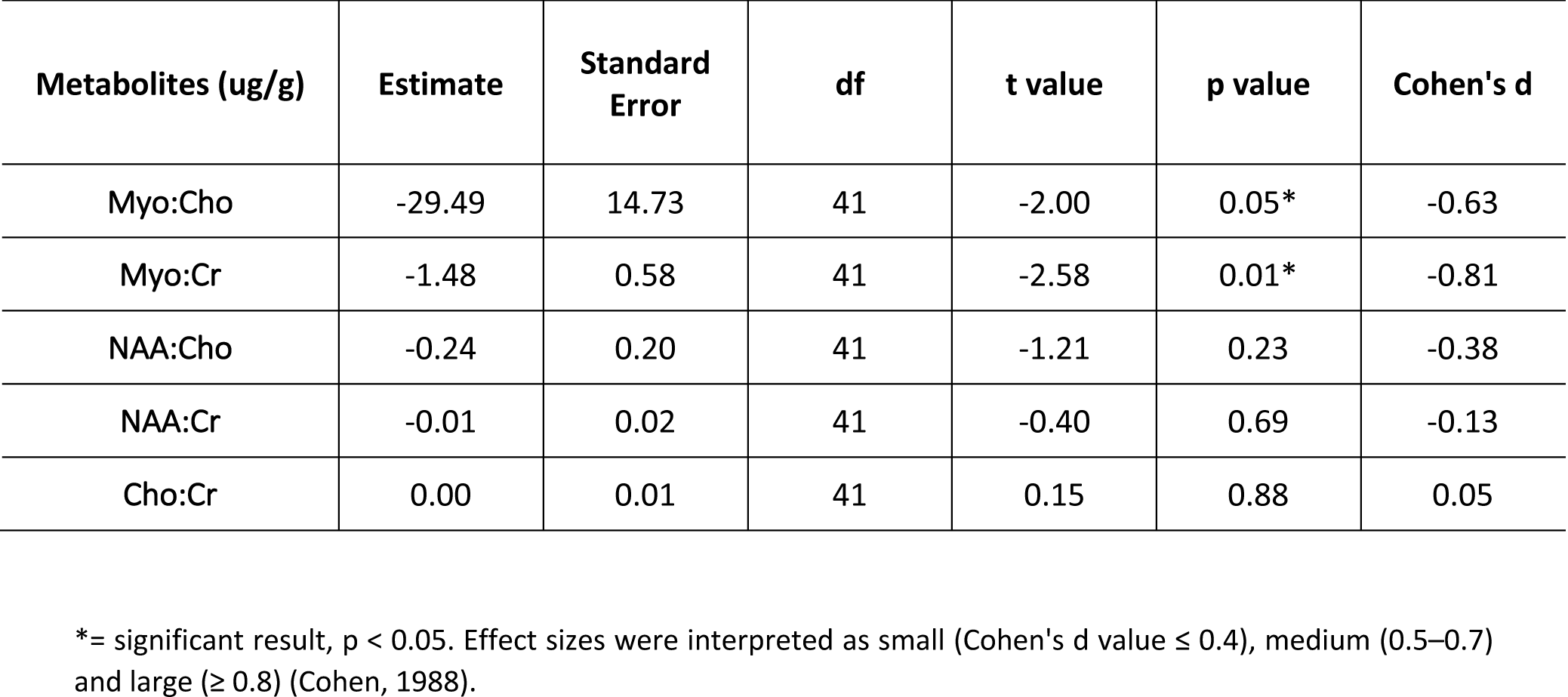
LME results, Cohen’s *d* values and effect sizes for the DS vs control cohort mass spectrometry ratios, as calculated from the absolute values

| Metabolites (ug/g) | Estimate | Standard Error | df | t value | p value | Cohen's d |
| --- | --- | --- | --- | --- | --- | --- |
| Myo:Cho | -29.49 | 14.73 | 41 | -2.00 | 0.05* | -0.63 |
| Myo:Cr | -1.48 | 0.58 | 41 | -2.58 | 0.01* | -0.81 |
| NAA:Cho | -0.24 | 0.20 | 41 | -1.21 | 0.23 | -0.38 |
| NAA:Cr | -0.01 | 0.02 | 41 | -0.40 | 0.69 | -0.13 |
| Cho:Cr | 0.00 | 0.01 | 41 | 0.15 | 0.88 | 0.05 |

\*= significant result, $p < 0.05$ . Effect sizes were interpreted as small (Cohen's *d* value $\leq 0.4$ ), medium (0.5–0.7) and large ( $\geq 0.8$ ) (Cohen, 1988).

## References

Acevedo, L. D., Holloway, H. W., Rapoport, S. I. and Shetty, H. U. (1997) ‘Application of stable isotope tracer combined with mass spectrometric detection for studying myo-inositol uptake by cultured neurons from fetal mouse: effect of trisomy 16.’, Journal of mass spectrometry : JMS, 32(4), pp. 395–400.

Baburamani, A. A., Patkee, P. A., Arichi, T. and Rutherford, M. A. (2019) ‘New approaches to studying early brain development in Down syndrome’, Developmental Medicine & Child Neurology, pp. 1–13.

Battin, M. and Rutherford, M. A. (2002) ‘Magnetic resonance imaging of the brain in preterm infants: 24 weeks’ gestation to term’, in MRI of the neonatal brain. W.B. Saunders.

Beacher, F. et al. (2005) ‘Hippocampal Myo-inositol and Cognitive Ability in Adults With Down Syndrome’, Arch Gen Psychiatry, 62, pp. 136–1365.

Berry, G., Wang, Z. and Dreha, S. (1999) ‘In vivo brain myo-inositol levels in children with Down syndrome’, The Journal of Pediatrics, 135(1), pp. 94–97.

Cárdenas, A. M., Fernández-Olivares, P., Díaz-Franulic, I., González-Jamett, A. M., Shimahara, T., Segura-Aguilar, J., Caviedes, R. and Caviedes, P. (2017) ‘Knockdown of Myo-Inositol Transporter SMIT1 Normalizes Cholinergic and Glutamatergic Function in an Immortalized Cell Line Established from the Cerebral Cortex of a Trisomy 16 Fetal Mouse, an Animal Model of Human Trisomy 21 (Down Syndrome)’, Neurotoxicity Research. Springer New York LLC, 32(4), pp. 614–623.

Chang, L., Munsaka, S. M., Kraft-Terry, S. and Ernst, T. (2013) ‘Magnetic resonance spectroscopy to assess neuroinflammation and neuropathic pain’, Journal of Neuroimmune Pharmacology. NIH Public Access, pp. 576–593.

Cohen, J. (1988) Statistical power analysis for the behavioral sciences. 2nd editio. Hillsdale, NJ: Lawrence Earlbaum Associates.

Contestabile, A., Greco, B., Ghezzi, D., Tucci, V., Benfenati, F. and Gasparini, L. (2013) ‘Lithium rescues synaptic plasticity and memory in Down syndrome mice.’, The Journal of clinical investigation, 123(1), pp. 348–61.

Coskun, P. E. and Busciglio, J. (2012) ‘Oxidative Stress and Mitochondrial Dysfunction in Down’s Syndrome: Relevance to Aging and Dementia’, Current Gerontology and Geriatrics Research. Hindawi Publishing Corporation, pp. 1–7.

Dai, G., Yu, H., Kruse, M., Traynor-Kaplan, A. and Hille, B. (2016) ‘Osmoregulatory inositol transporter SMIT1 modulates electrical activity by adjusting PI(4,5)P2 levels’, Proceedings of the National Academy of Sciences of the United States of America. National Academy of Sciences, 113(23), pp. E3290–E3299.

Fenili, D., Weng, Y.-Q., Aubert, I., Nitz, M. and McLaurin, J. (2011) ‘Sodium/myo-Inositol Transporters: Substrate Transport Requirements and Regional Brain Expression in the TgCRND8 Mouse Model of Amyloid Pathology’, PLoS ONE. edited by C. V. Borlongan, 6(8), p. e24032.

Ferguson, K. J., Maclullich, A. M. J., Marshall, I., Deary, I. J., Starr, J. M., Seckl, J. R. and Wardlaw, J. M. (2002) ‘Magnetic resonance spectroscopy and cognitive function in healthy elderly men’, Brain, 125, pp. 2743–2749.

Fisher, S. K., Novak, J. E. and Agranoff, B. W. (2002) ‘Inositol and higher inositol phosphates in neural tissues: Homeostasis, metabolism and functional significance’, Journal of Neurochemistry, 82(4), pp. 736–754.

Girard, N., Fogliarini, C., Viola, A., Confort-Gouny, S. Fur, Y. Le, Viout, P., Chapon, F., Levrier, O. and Cozzone, P. (2006) ‘MRS of normal and impaired fetal brain development.’, European journal of radiology, 57(2), pp. 217–25.

Harris, J. L., Choi, I.-Y., Brooks, W. M., Barreto, G. E., Oliveira, C. and Arturo García-Horsman, J. (2015) ‘Probing astrocyte metabolism in vivo: proton magnetic resonance spectroscopy in the injured and aging brain’, Frontiers in Aging Neuroscience, 7.

Holub, B. J. (1986) Metabolism and function of myo-inositol and inositol phospholipids, Ann. Rev. Nutr.

Huang, W., Alexander, G. E., Daly, E. M., Shetty, H. U., Krasuski, J. S., Rapoport, S. I. and Schapiro, M. B. (1999) ‘High Brain myo-Inositol Levels in the Predementia Phase of Alzheimer’s Disease in Adults With Down’s Syndrome: A 1 H MRS Study’, Am J Psychiatry, 156, pp. 1879–1886.

Hughes, E. J. et al. (2017) ‘A dedicated neonatal brain imaging system’, Magnetic Resonance in Medicine, 78, pp. 794–804.

Jansen, J. F. A., Backes, W. H., Nicolay, K. and Kooi, M. E. (2006) ‘1H MR Spectroscopy of the Brain: Absolute Quantification of Metabolites’, Radiology, 240(2), pp. 318–332.

Karmiloff-Smith, A. et al. (2016) ‘The importance of understanding individual differences in Down syndrome’, F1000Research, 5, p. 389.

Kato, T., Nishina, M., Matsushita, K., Hori, E., Mito, T. and Takashima, S. (1997) ‘Neuronal maturation and N-acetyl-L-aspartic acid development in human fetal and child brains’, Brain & Development, 19, pp. 131–133.

Kim, D. H., Barkovich, A. J. and Vigneron, D. B. (2006) ‘Short Echo Time MR Spectroscopic Imaging for Neonatal Pediatric Imaging’, American Journal of Neuroradiology, 27, pp. 1370–1372.

Kok, R. D., van den Bergh, A. J., Heerschap, A., Niljand, R. and van den Berg, P. P. (2001) ‘Metabolic information from the human fetal brain obtained with proton magnetic resonance spectroscopy’, American Journal of Obstetrics and Gynecology, 185(5), pp. 1011–1015.

Kok, R. D., van den Berg, P. P., van den Bergh, A. J., Nijland, R. and Heerschap, A. (2002) ‘Maturation of the human fetal brain as observed by1H MR spectroscopy’, Magnetic Resonance in Medicine, 48(4), pp. 611–616.

Lally, P. J. et al. (2019) ‘Magnetic resonance spectroscopy assessment of brain injury after moderate hypothermia in neonatal encephalopathy: a prospective multicentre cohort study’, Lancet Neurology, 18, pp. 35–45.

Lin, A. L. et al.(2016) ‘1H-MRS metabolites in adults with Down syndrome: Effects of dementia’, NeuroImage: Clinical, 11, pp. 728–735.

Murphy, D. (2011) ‘Genetically determined brain abnormalities in Down syndrome – towards a treatment?: A single blind placebo controlled trial of lithium carbonate in Down Syndrome’, (February).

Öz, G. et al. (2014) ‘Clinical Proton MR Spectroscopy in Central Nervous System Disorders’, Radiology, 270(3), pp. 658–679.

Patkee, P. A. et al. (2020) ‘Early alterations in cortical and cerebellar regional brain growth in Down Syndrome: An in vivo fetal and neonatal MRI assessment’, NeuroImage: Clinical, 25, p. 102139.

Rachidi, M. and Lopes, C. (2007) ‘Mental retardation in Down syndrome: from gene dosage imbalance to molecular and cellular mechanisms.’, Neuroscience research, 59(4), pp. 349–69.

Rafii, M. S., Skotko, B. G., Mcdonough, M. E., Pulsifer, M., Evans, C., Doran, E., Muranevici, G., Kesslak, P. and Abushakra, S. (2018) ‘A Randomized, Double-Blind, Placebo-Controlled, Phase 2 Study of Oral ELND005 (scyllo-Inositol) in Young Adults with Down Syndrome without Dementia’, J Alzheimers Dis, 58(2), pp. 401–411.

Rosenthal, R. and Rosnow, R. (2008) Essentials of Behavioral Research: Methods and Data Analysis. Third edit. New York: McGraw-Hill.

Santamaria, A., Corrado, F., Interdonato, M. L., Baviera, G., Carlomagno, G., Cavalli, P., Unfer, V. and D’Anna, R. (2014) ‘Myo-inositol in Down syndrome amniotic fluid. A case-control study’, Prenatal Diagnosis, pp. 917–918.

Shetty, H. U., Schapiro, M. B., Holloway, H. W. and Rapoport, S. (1995) ‘Polyol Profiles in Down Syndrome’, The Journal of clinical investigation, 95(February), pp. 542–546.

Shonk, T. and Ross, B. D. (1995) ‘Role of increased cerebral myo-inositol in the dementia of Down syndrome.’, Magnetic resonance in medicine : official journal of the Society of Magnetic Resonance in Medicine / Society of Magnetic Resonance in Medicine, 33(6), pp. 858–61.

Siger, M., Schuff, N., Zhu, X., Miller, B. L. and Weiner, M. W. (2009) ‘Regional Myo-inositol Concentration in Mild Cognitive Impairment Using 1H Magnetic Resonance Spectroscopic Imaging’, Alzheimer Dis Assoc Disord, 23(1), pp. 57–62.

Śmigielska-Kuzia, J., Boćkowski, L., Sobaniec, W., Kułak, W. and Sendrowski, K. (2010) ‘Amino acid metabolic processes in the temporal lobes assessed by proton magnetic resonance spectroscopy (1H MRS) in children with Down syndrome.’, Pharmacological reports, 62(6), pp. 1070–7.

Tarui, T. et al. (2019) ‘Quantitative MRI Analyses of Regional Brain Growth in Living Fetuses with Down Syndrome’, Cerebral Cortex. Oxford University Press (OUP), 00(00), pp. 1–9.

Tomiyasu, M., Aida, N., Endo, M., Shibasaki, J., Nozawa, K., Shimizu, E., Tsuji, H. and Obata, T. (2013) ‘Neonatal Brain Metabolite Concentrations: An In Vivo Magnetic Resonance Spectroscopy Study with a Clinical MR System at 3 Tesla’, PLoS ONE, 8(11), pp. 1–7.

Wiseman, F., Al-Janabi, T., Karmiloff-Smith, A., Nizetic, D., Tybulewicz, V., Fisher, E. and Strydom, A. (2015) ‘A genetic cause of Alzheimer disease: mechanistic insights from Down syndrome’, Nature reviews Neuroscience, 16(9), pp. 564–574.

Wisniewski, K. and Schmidt-Sidor, B. (1989) ‘Postnatal delay of myelin formation in brains from Down syndrome infants and children’, Clinical neuropathology, 8(2), pp. 55–62.

Yuan, J. et al. (2018) ‘Lithium Treatment Is Safe in Children With Intellectual Disability’, Front. Mol. Neurosci, 11, p. 425.

